# Combining xenium *in situ* spatial transcriptomics and imaging mass cytometry on a single tissue section

**DOI:** 10.64898/2026.02.18.700929

**Authors:** Ruth O Allen, Farhan Ameen, Emily Duchini, Thomas Ashhurst, Rachael Ireland, Jordan W Conway, Xinyu Bai, Angela M Hong, Angela L Ferguson, Ellis Patrick, Umaimainthan Palendira

## Abstract

Spatial imaging technologies provide an expansive view of tissue microenvironments through high-plex profiling of protein and molecular targets *in situ*. Imaging mass cytometry (IMC; Standard BioTools) is a trusted method for defining immune phenotypes based on up to 40 protein targets, whilst Xenium *in situ* spatial transcriptomics (Xenium; 10x Genomics) is an emerging platform that can measure up to 5000 mRNA markers simultaneously. Although these platforms can reveal valuable insights on their own, there is an increasing need to analyse samples using a multi-omics approach to further our understanding of complex biological processes. To address this, we have assessed a novel dual-platform workflow that combines Xenium and IMC on a single formalin-fixed paraffin-embedded tissue section to enable the spatial profiling of both mRNA and protein targets at single-cell resolution. The feasibility of the workflow was determined by comparing the staining quality of IMC performed after Xenium to that of IMC performed alone on an adjacent tissue section, confirming that Xenium has little to no negative impact on subsequent IMC protein staining. Although the location of transcripts picked up by Xenium correlated with the corresponding proteins picked up by IMC at a global scale, discrepancies between the two technologies were apparent at the single-cell level. This is to be expected, as biologically transcript expression does not always correlate with protein, and both platforms have their own technical limitations. However, when we analyse T cells identified by both technologies, as opposed to T cells identified by Xenium or IMC alone, it produces the most biologically meaningful results at both the transcript and protein level for specific T cell markers. These results highlight how integration of the two platforms, identifying the presence of both RNA and protein, can foster a more comprehensive view of cellular landscapes and provide a greater depth of functional capabilities and cellular interactions.

## 1 Introduction

Spatial omics technologies provide an expansive view of tissue microenvironments through high-plex profiling of an increasing number of protein and molecular targets *in situ* (Bressan et al., 2023). In particular, imaging mass cytometry (IMC^TM^; Standard BioTools) is a trusted method for defining detailed cell phenotypes and mapping cell-cell interactions based on up to 40 protein markers (Glasson et al., 2023; Kakade et al., 2021), whilst Xenium^TM^ *in situ* spatial transcriptomics (Xenium; 10x Genomics) is an emerging platform that can measure up to 5000 mRNA targets simultaneously (Azevedo, 2024). Although these single-cell resolved technologies reveal valuable insights on their own (Ferguson et al., 2022; Janesick et al., 2023), there is an increasing need to analyse tissue samples using a multi-omics approach to achieve a more holistic and accurate view of cellular landscapes and further our understanding of complex biological disease processes. Multi-omic workflows on same-tissue sections have been applied in other studies to address this, with most focusing on combining transcriptomic and proteomic information although not necessarily at single cell resolution (Ben-Chetrit et al., 2023; Enninful et al., 2026; Guilliams et al., 2022; Hendriks et al., 2025; Huynh et al., 2025; Liu et al., 2020; Liu et al., 2023; Merritt et al., 2020; Tran et al., 2025; Vickovic et al., 2022; Yeo et al., 2024). Whilst protein expression is more dependable for phenotypic identification and reflective of cellular states and functions, the number of targets that can be simultaneously profiled at the single-cell level is limited to around 100 and currently can only be achieved on one platform (He et al., 2022). Therefore, it is advantageous to supplement proteomic insights with same-tissue high-plex transcriptomics to uncover more comprehensive insights into cellular and molecular pathways, ultimately enabling the more meaningful determination of cell phenotype, function and potential. Although methods like cellular indexing of transcriptomes and epitopes by sequencing (CITE-seq) were developed to address the need of both proteomic and transcriptomic information from the same cell, these single-cell approaches do not retain the spatial context of tissue samples (Stoeckius et al., 2017).

Currently, the CosMx^TM^ Spatial Molecular Imager and Xenium Protein assay are the only spatial solutions on the market that enable simultaneous mRNA and protein profiling at single-cell resolution (10x Genomics, 2026; He et al., 2022). However, both technologies offer only validated subpanels for proteomic analysis, preventing users from building a panel tailored to their tissue and markers of interest. This can be overcome by combining two separate platforms on a single tissue section, whereby the proteomic platform enables customised protein detection. This has been demonstrated by previous workflows that have combined Xenium with fluorescence-based platforms, either PhenoCycler^TM^-Fusion (PCF; Akoya Biosciences) (Huynh et al., 2025) or COMET^TM^ (Lunaphore) (Rademacher et al., 2025; Tran et al., 2025), signalling the possibility of performing additional high-plex analysis on processed Xenium slides. IMC is an alternative proteomic imaging technology and has previously been used to validate or enrich findings from spatial transcriptomics on adjacent tissue sections (Bell et al., 2024; Liu et al., 2024; Ma et al., 2025). This technology is characterised by the use of metal-tagged antibodies for protein detection, contributing to low background signal, no autofluorescence and a short staining protocol (Kakade et al., 2021). There is growing interest to combine spatial transcriptomic and proteomic information by performing Xenium and IMC on a single tissue slide (Khakpoor et al., 2025; Marlin et al., 2025). However, to date, there are no publications that have assessed the feasibility of this workflow or have comprehensively demonstrated integration of the two technologies.

Here, we describe the novel dual-platform workflow that combines Xenium and IMC on a single formalin-fixed paraffin-embedded (FFPE) tissue section to enable spatial profiling of both mRNA and protein at single-cell resolution. The ability to perform this workflow on the same slide is advantageous for avoiding problems that arise when performing experiments on adjacent tissue sections, including subsequent spatial differences which result in slight variations in cellular composition between sections. While this study was performed on a limited number of samples, the primary aim was to establish technical feasibility and analytical concordance of a same-section dual-platform workflow rather than draw population-level biological conclusions. This method has potential to contribute to more accurate characterisation of cell populations and spatially informed interaction analysis, thus providing greater depth of functional capabilities and cellular interactions. Furthermore, this workflow can also act as a validation tool for both technologies whilst addressing their respective limitations.

## 2 Methods

### 2.1 Sample Collection

This study was performed on a FFPE tissue microarray (TMA) containing 1 mm^2^ cores of melanoma cells from an involved lymph-node. The samples in this study were part of a phase 3 randomised controlled trial for adjuvant lymph-node field radiotherapy versus observation only in patients with melanoma and were not subjected to neoadjuvant systemic therapy [ANZMTG 01.02/TROG 02.01] (Henderson et al., 2015, Burmeister et al., 2012).

### 2.2 Sample Preparation

5 µm tissue sections were mounted on Xenium slides following 10X Genomics protocols (Document CG000578, Rev C). The TMA sample block was scored into two rectangular areas that separated in the water bath after sectioning to accommodate the larger size of the full TMA area to the restricted sample area of the Xenium slides (12 mm x 24 mm). The two TMA sample sections were mounted onto two separate Xenium slides, labelled Slide A and Slide B, and left to dry at room temperature (RT). The slides were then placed on the Xenium Cassette Adaptor (10X Genomics) in a C1000 Thermal Cycler (Bio-Rad) at 42°C for 3 hours prior to being stored in a falcon tube containing a desiccator overnight.

### 2.3 Xenium *In Situ* Spatial Transcriptomics

#### 2.3.1 Tissue Deparaffinization and Decrosslinking

Deparaffinisation and decrosslinking were performed according to 10X Genomics protocols (Document CG000580, Rev C). Slides were baked in the thermal cycler at 60°C for 120 minutes then left to cool on the bench for 10 minutes. The tissue was deparaffinized by submerging in two jars containing xylenes (Sigma Aldrich) consecutively for 10 minutes each. Slides were then rehydrated through a graded series of ethanol solutions (Sigma) for 3 minutes each at the following concentrations: 100%, 100%, 96%, 96%, 70%, before a wash in nuclease-free water (Thermo Fisher Scientific) for 20 seconds. Slides were then placed within custom Xenium Cassettes to create a physical seal around the tissue area. Slides were first washed with 500 µL of PBS (Invitrogen), followed by 500 µL of Decrosslinking Buffer (Xenium Slides & Sample Prep Reagents, 10X Genomics, PN-1000460) before being placed in the thermal cycler with the following settings: 22°C ‘hold’, 80°C for 30 minutes, 22°C for 10 minutes, 22°C ‘hold’.

#### 2.3.2 Xenium *In Situ* Gene Expression Assay

Samples were analysed using the Xenium Human Multi-Tissue and Cancer Expression Panel (10X Genomics, PN-1000626) containing 377 gene target probes purchased in February 2024. Tissue hybridisation, ligation, amplification and autofluorescence quenching was performed according to 10X Genomics protocols (Document CG000582, Rev E) using Xenium Slides & Sample Prep Reagents (10X Genomics, PN-1000460). After decrosslinking, slides were washed with 500 µL of PBS-T (PBS containing 0.05% Tween 20) three times for 1 minute each. 500 µL of Probe Hybridisation Mix (10X Genomics) was added to samples before being placed in the thermal cycler with the following settings: 50°C ‘hold’, 16-24 hour/overnight at 50°C, followed by a 50°C ‘hold’. Slides were then washed with 500 µL of PBS-T twice before 500 µL of Xenium Post Hybridisation Wash Buffer (10X Genomics) was applied to samples. Slides were then placed on the thermocycler for 30 minutes at 37°C and subsequently washed with 500 µL of PBS-T before 500 µL of Ligation Mix (10X Genomics) was added to samples. Slides were then placed within the thermal cycler for 120 minutes at 37°C. Following ligation of the gene probes, slides were washed with PBS-T three times before 500 µL of Amplification Master Mix (10X Genomics) was applied to samples and placed in the thermal cycler for 120 minutes at 30°C. Slides were then washed with 500 µL of TE Buffer (Thermo Fisher Scientific) three times, then another three times with 1000 µL of PBS. 500 µL of Diluted Reducing Agent B (10X Genomics) was applied to samples and incubated for 10 minutes at RT. Samples were then washed with 1000 µL of 70% ethanol, followed by two further washes with 100% ethanol. 500 µL of Autofluorescence Solution (10X Genomics) was applied to samples and incubated for 10 minutes at RT in the dark. Samples were then washed with 1000 µL of 100% ethanol three times, with no further solution added before being placed in the thermal cycler and dried for 5 minutes at 37°C. Slides were then rehydrated with 1000 µL of PBS for 1 minute, followed by 1000 µL of PBS-T for 2 minutes in the dark. 500 µL of Xenium Nuclei Staining Buffer (10X Genomics) was applied to samples and incubated for 1 minute at RT in the dark. Slides were then washed with 1000 µL of PBS-T three times for 1 minute in the dark. 1000 µL of PSB-T was then added to slides before loading into the Xenium Analyzer (10X Genomics) following 10X Genomics protocols (Document CG000584, Rev D), for automated cyclic imaging and on-instrument Xenium Onboard Analysis (XOA, v1.9).

#### 2.3.3 Xenium Preliminary Data Analysis

The Xenium Experiment output file was imported into Xenium Explorer 3 (10X Genomics, v3.2.0) for visualisation and to produce high resolution PNG representative images. The cell segmentation mask generated by the XOA pipeline was used for preliminary analysis, where cell boundaries were individually identified by nuclear expansion of 15 µm or until another cell boundary was encountered (10x Genomics, 2025). Cell transcript expression and area was exported for each TMA sample core as a comma separated value (CSV) file.

### 2.4 Imaging Mass Cytometry (IMC) Spatial Proteomics

#### 2.4.1 IMC Post-Xenium

Following acquisition on the Xenium Analyzer, Slide A was stained for IMC using a 40-plex metal-tagged antibody panel (Table 1). All antibodies were conjugated, titrated for optimal concentration, and prepared as master mixes by Sydney Cytometry. The tissue section was washed in TRIS-Buffered Saline with Tween-20 (TBS-T) (0.1 M Tris HCI, 0.15 M NaCl, 0.05% Tween-20 made up in deionised H2O, adjusted to pH 7.5) twice for 2 minutes each. Antigen retrieval was performed by submerging the tissue section in pH 9.0 antigen retrieval buffer (10 mM Tris Base, 1 mM EDTA, 0.05% Tween-20, adjusted to pH9) and microwaving at full power (100% on a 700W output microwave) until boiling, followed by an additional 15 minutes maintained at boiling (low/20% power). The tissue section was cooled to RT before two subsequent washes in TBS-T and Dulbecco’s Phosphate-Buffered Saline (DPBS) (Gibco) for 10 minutes each. The tissue section was then blocked with 1X Antibody Diluent/Block solution (Akoya Biosciences) for 45 minutes at 37°C in a hydration chamber. The tissue section was then stained with anti-human SOX10 (clone BC34, Biocare Medical) diluted 1:100 in 1X Antibody Diluent/Block for 45 minutes at RT. The tissue section was then washed twice in TBS-T before staining with AF647/Cy5 anti-mouse (polyclonal, Jackson ImmunoResearch Laboratories) diluted 1:100 in DPBS for 30 minutes at RT. The tissue section was washed twice in TBS-T before incubation with the metal-tagged antibody master mix (Table 1) at 4°C overnight in a hydration chamber. The tissue section was then washed in 0.1% Triton-X (Sigma) diluted in DPBS twice for 8 minutes each, followed by two washes in DPBS for 8 minutes each. The tissue section was then stained with Cell-ID Ir-Intercalator (Standard BioTools) diluted 1:400 in DPBS for 30 minutes at RT. The tissue section was then washed with deionised water for 5 minutes then let to air dry at RT. Samples were acquired using the Hyperion Imaging System (Standard BioTools) with pulsed laser ablation performed at 200 Hz.

**Table 1:** Imaging mass cytometry antibody panel.

| **Metal** | **Marker** | **Clone** | **Company** | **Concentration**<br>**(µg/mL)** |
| --- | --- | --- | --- | --- |
| 89Y | alpha-SMA | 1A4 | R&D Systems | 1.00 |
| 113In | Vimentin | D21H3 | Cell Signalling Technology | 2.00 |
| 115In | CD11c | D3V1E | Cell Signalling Technology | 3.99 |
| 139La | pan-Cytokeratin | SPM115/SPM116 | R&D Systems | 1.00 |
| 141Pr | CD20 | H1 | Becton Dickinson | 4.00 |
| 142Nd | HH3 | polyclonal | Novus Biologicals | 4.02 |
| 143Nd | CD45RA | HI100 | Becton Dickinson | 4.00 |
| 144Nd | Myeloperoxidase | E1E71 | Cell Signalling Technology | 7.99 |
| 145Nd | CD103 (ITGAE) | EPR4166(2) | Abcam | 3.01 |
| 146Nd | CD8a | D8A8Y | Cell Signalling Technology | 3.52 |
| 147Sm | Podoplanin | polyclonal | R&D Systems | 2.00 |
| 148Nd | CD16 | EPR16784 | Abcam | 8.00 |
| 149Sm | CADM1 | polyclonal | Invitrogen | 5.00 |
| 150Nd | IDO | D5J4E | Cell Signalling Technology | 8.00 |
| 151Eu | CD274 (PD-L1) | E1L3N | Cell Signalling Technology | 2.00 |
| 152Sm | CD13 | 498001 | R&D Systems | 4.01 |
| 153Eu | CD68 | KP1 | Biolegend | 2.00 |
| 154Sm | VISTA | D1L2G | Cell Signalling Technology | 7.98 |
| 155Gd | CD31 | MEC13.3 | Novus Bio | 4.00 |
| 156Gd | CD183 (CXCR3) | 49801 | Abcam | 1.00 |
| 158Gd | T-bet | D6N8B | Cell Signalling Technology | 6.00 |
| 159Tb | CD197 (CCR7) | G043H7 | Biolegend | 8.01 |
| 160Gd | CD14 | EPR3653 | Abcam | 2.00 |
| 161Dy | FXIIIa | polyclonal | Affinity Biologicals | 7.68 |
| 162Dy | FoxP3 | polyclonal | R&D Systems | 4.00 |
| 163Dy | CD279 (PD1) | EPR4877(2) | Abcam | 2.00 |
| 164Dy | SIRPa | D6I3M | Cell Signalling Technology | 3.99 |
| 165Ho | OX40 | E9U70 | Cell Signalling Technology | 3.99 |
| 166Er | CD45 | D9M8I | Cell Signalling Technology | 1.99 |
| 167Er | CD66 | YTH71.3 | Abcam | 1.00 |
| 168Er | Ki67 | B56 | Becton Dickinson | 1.98 |
| 169Tm | anti-AF647/Cy5 | CY5-15 | Sigma | 1.00 |
| 170Er | CD3 | polyclonal | Abcam | 4.61 |
| 171Yb | Granzyme B | EPR8260 | Abcam | 8.03 |
| 172Yb | CD206 (MMR) | 685645 | R&D Systems | 4.00 |
| 173Yb | CD4 | EPR6855 | Abcam | 4.00 |
| 174Yb | HLA-DR | EPR3692 | Abcam | 1.00 |
| 175Lu | ICOS | D1K2T | Cell Signalling Technology | 10.02 |
| 176Yb | CD56 | EPR2566 | Abcam | 4.00 |
| 209Bi | DC SIGN | polyclonal | Abcam | 1.00 |

#### 2.4.2 IMC Alone

For sections analysed by IMC alone, slides were first baked at 60°C for 60 minutes before deparaffinization in xylenes for 10 minutes. Slides were then rehydrated through a graded series of ethanol solutions for 5 minutes each at the following concentrations: 100%, 90%, 70%, 50%. Slides were washed in TBS-T twice for 2 minutes each prior to antigen retrieval. Antigen retrieval and all subsequent staining steps were followed as above (2.4.1 IMC Post-Xenium).

#### 2.4.3 IMC Preliminary Data Analysis

The mass cytometry data (mcd) file for each IMC experiment was imported into MCD viewer (Standard BioTools, v1.0.560.6) for visualisation and to produce representative tagged image file format (TIFF) images. Each TMA sample core was exported as single-marker TIFFs for cell segmentation using Cell Profiler (v4.2.8; Stirling et al., 2021), where cells were individually identified based on nuclear staining (DNA-Ir). Each TMA sample was then exported as a CSV file that contained each segmented cell as a data point with *x,y* coordinates and marker expression. The csv files were imported into FlowJo (BD Bioscience, v10.10.0) where manual gating of cell populations was performed. The area of each sample was calculated using manual annotations of tissue sections with the selection tool in Fiji (v1.54p), which was subsequently used to calculate the density of cell populations.

### 2.5 Hematoxylin and Eosin Staining

Following acquisition on the Xenium Analyzer, Slide B was stained for H&E. The slide was immersed in Milli-Q dH2O three times for 1 minute each, followed by Hematoxylin (Sigma-Aldrich) for 3 minutes. The slide was washed again in Milli-Q dH2O three times prior to staining with Blueing Solution (Hurst Scientific) for 1 minute, before being washed in Milli-Q dH2O for 1 minute. The slide was then placed in a graded series of ethanol solutions for 1 minute each at the following concentrations: 50%, 70%, 95%, 95%. The slide was then counterstained in Eosin (Sigma-Aldrich) for 45 seconds, followed by immersion in xylenes for 2 minutes. The slide was then mounted using ProLong Gold Antifade Mountant (Invitrogen) and stored at RT in the dark. Imaging was performed on the ZEISS Axioscan.

### 2.6. Integrated Multi-omic Data Analysis

#### 2.6.1 Registration of the Xenium and IMC Datasets

The Xenium and IMC datasets were first aligned by registering the corresponding nuclear images using the Tissuealign analysis module in Visiopharm (v2025.02.1). The *morphology_mip* TIFF file from the XOA outputs was used to visualise the Xenium DAPI image. The IMC single-colour TIFF images exported per sample were combined into a hyper stack using Fiji. The Xenium *morphology_mip* TIFF and IMC hyper stack OME-TIFF were imported into the Visiopharm database. The two images were linked by first selecting the Xenium *morphology_mip* TIFF to make it the reference image, then the IMC hyper stack OME-TIFF was selected so that the two images could be viewed side by side. 320 pins were manually placed on visually distinct locations in the Xenium DAPI image and corresponding IMC DNA-Ir image to achieve good registration of the IMC image to the Xenium image. The aligned IMC image was exported as a TIFF file.

#### 2.6.2 Segmentation

The Xenium image was read into R and converted to a ‘SingleCellExperiment’ object using the ‘readXenium’ and ‘countMolecules’ functions in the MoleculeExperiment R package (Peters Couto et al., 2025). The aligned IMC image was scaled up by a factor of 0.2125 and flipped horizontally to match the pixel coordinates of the Xenium mask. The Xenium cell boundaries were converted into polygons using the ‘st_sf’ function from the sf R package (Pebesma and Bivand, 2023; Pebesma, 2018). The resulting polygons were converted to a raster mask using the fasterize R package (Ross et al., 2025). Finally, the aligned IMC image was segmented using the ‘measureObjects’ function in the cytomapper R package (Eling et al., 2020) (Figure 3D).

#### 2.6.3 Normalisation

The protein intensities in the IMC image were normalised using the ‘normalizeCells’ function from the simpleSeg R package (Canete et al., 2025), applying PC1, trim99 and minMax normalisation. The following background and nuclei specific markers were omitted for normalising: 80ArAr, 104Pd, 189Os, 190Os, 208Pb, HH3, DNA1, DNA2. In the Xenium image, any cell containing 0 transcripts were filtered out. Transcripts counts were normalised using the ‘logNormCounts’ function in the scater R package (McCarthy et al., 2017).

#### 2.6.4 Cell type annotation

B cells and T cells were annotated using two approaches. In the first approach, cells were identified based solely on the presence or absence of a marker. Using the normalised protein intensities in the IMC image, a cell was labelled as a T cell if CD3 > 0.4 and CD20 < 0.065, whilst a B cell was identified if CD20 > 0.065 and CD3 < 0.4. These numbers were determined by manual gating and thresholding in FlowJo. Using the normalised transcript values from Xenium, cells were labelled as T cells or B cells if they contained at least 1 transcript of *CD3E* or *MS4A1* respectively.

In the second approach, cells were identified based on semi-supervised clustering. Using the normalised protein intensities, the cells were clustered into 10 clusters using the FuseSOM R package (Willie et al., 2023). The following cell type specific markers were used to cluster cell types: CD11c, CD68, CD20, CD8a, Podoplanin, CD14, FX111A, CD45, CD66, CD3, SOX10, HLA-DR, CD4, and CD31. The average expression of these markers was standardised and visualised for each cluster using the ‘plotGroupedHeatmap’ function from scater (McCarthy et al., 2017). Using the normalised transcript values, negative control probes were filtered out then cells were clustered using the clustSIGNAL R package (Panwar et al., 2025), using default parameters. The top 5 differentially expressed genes were identified for each cluster using the ‘findMarkers’ function from the scran R package (Lun et al, 2016). The average expression of these transcripts was standardised and visualised for each cluster using the ‘plotGroupedHeatmap’ from scater (McCarthy et al., 2017). Cell type annotation for this second approach was performed manually based on the resulting heatmaps of cluster marker expression (Figure 4A, B).

#### 2.6.5 Constructing concordance masks

Concordance masks were constructed using the cell type annotations from the two separate approaches. For the first approach, T cells were labelled as “identified by both Xenium and IMC” if the cell was positive for CD3E (*CD3E*) in both transcript and protein, and “identified by IMC” or “identified by Xenium” if positive for CD3E in just one modality. B cells were labelled the same way, instead using CD20 and *MS4A1* for the proteins and transcripts respectively. Similarly for the second approach, T cells and B cells were labelled as “identified by both Xenium and IMC” if the cell was manually annotated as a T cell or B cell in both datasets, and “identified by IMC” or “identified by Xenium” if manually annotated as a T cell or B cell in just one modality. Finally, the ‘plotCells’ function from cytomapper (Eling et al., 2020) was used to visualise the marker based and cell type annotation-based concordance masks (Figure 4C, D).

#### 2.6.6 Region analysis

Regions were identified using the lisaClust R package (Patrick et al., 2023) on IMC typed cells with k = 4 and were manually annotated based on the distribution of cell types within each region (Figure 4I). T cells were then labelled as either inside or outside the tumour region (Figure 4J) and further stratified by whether they were identified as T cells by both technologies or by a single modality alone. For each stratification, the average expression of key T cell activation markers (CD3, Ki67, PD-1, Lag3, GranzymeB, FoxP3) and their corresponding transcripts (*CD3E, MKI67, PDCD1, LAG3, GZMB, FOXP3*) was visualised using the ‘plotGroupedHeatmap’ function in scater (McCarthy et al., 2017) (Figure 4K).

## 3.0 Results

### 3.1 A novel workflow combining Xenium spatial transcriptomics and IMC proteomics on a single tissue section

The novel dual-platform workflow was performed on two FFPE TMA tissue sections, labelled Slide A and Slide B. Each TMA section contained 1 mm^2^ tissue cores from melanoma lymph node tumour samples. Spatial transcriptomics was first performed using the Xenium platform in accordance with 10X Genomics protocols (Figure 1A). Briefly, the samples (Slide A and B) underwent probe ligation and rolling circle amplification prior to being loaded into the Xenium Analyzer, where cyclic probe addition, imaging, and removal was performed automatically to spatially map 377 mRNA targets across the tissue sections. After the run was complete, Slide B underwent H&E staining to assess the impact of the Xenium workflow on the tissue integrity (Figure 1C), whilst Slide A underwent IMC multiplexed protein imaging (Figure 1B). The section (Slide A) was stained with metal-tagged monoclonal antibodies prior to acquisition using the Hyperion Imaging System, where laser ablation generated plumes of metal ions for detection by time-of-flight mass cytometry to profile 40 protein targets in the tissue at a sub-cellular resolution.

**Figure 1.**
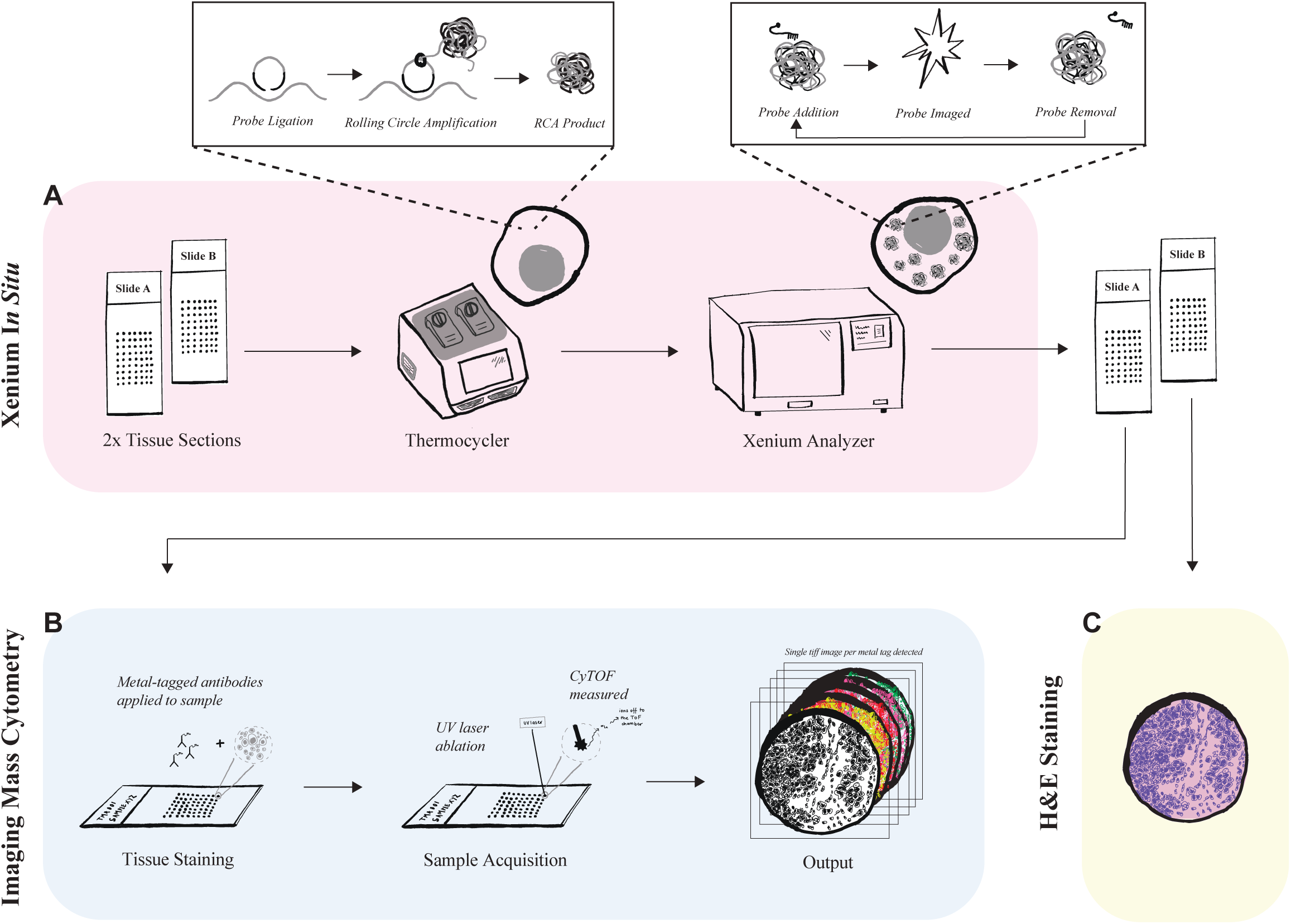
Combined Xenium and IMC experimental workflow performed on a single FFPE tissue section. **(A)** Two FFPE TMA tissue sections, labelled Slide A and Slide B, firstly underwent Xenium *In Situ* spatial transcriptomics. Briefly, slides underwent probe ligation and rolling circle amplification (RCA) in a thermocycler, prior to being loaded into the Xenium Analyzer where probes were added, imaged and removed cyclically. (**B**) Following Xenium, Slide A underwent high-dimensional protein evaluation with Imaging Mass Cytometry. The tissue section (Slide A) was stained using metal-conjugated antibodies prior to acquisition using the Hyperion Imaging System, where UV laser ablation generated plumes of metal ions for detection by time-of-flight mass cytometry (CyTOF), ultimately generating single TIFF images for each protein target. (**C**) The second tissue section, Slide B, underwent H&E staining following Xenium.

### 3.2 Xenium has minimal impact on tissue quality and subsequent IMC proteomic staining

One of the major concerns of combining spatial transcriptomics and targeted proteomics on the same section is the possible loss of tissue from the section and reduced quality of the subsequent protein staining. We therefore implemented multiple quality control measures to determine the feasibility of performing IMC after Xenium on the same tissue section. First, we performed H&E staining of Slide B after Xenium and compared it to the DNA-DAPI image obtained by the Xenium Analyser at the start of the imaging cycle, to visually inspect the impact of the Xenium workflow on the integrity of the tissue (Figure 2A). H&E staining revealed that the tissue is well maintained after Xenium with minimal tissue loss, and is therefore suitable for successive IMC staining. This was supported by the high visual concordance of the Xenium DNA-DAPI and the corresponding IMC DNA-Ir images, obtained by the Xenium Analyzer and Hyperion Imaging System respectively, for the samples on Slide A that underwent the dual-platform workflow (Figure 2B). This concordance, together with the tissue being able to withstand two heat-induced antigen retrievals in the IMC staining protocol (Slide A), demonstrates that Xenium has minimal impact on tissue integrity.

**Figure 2.**
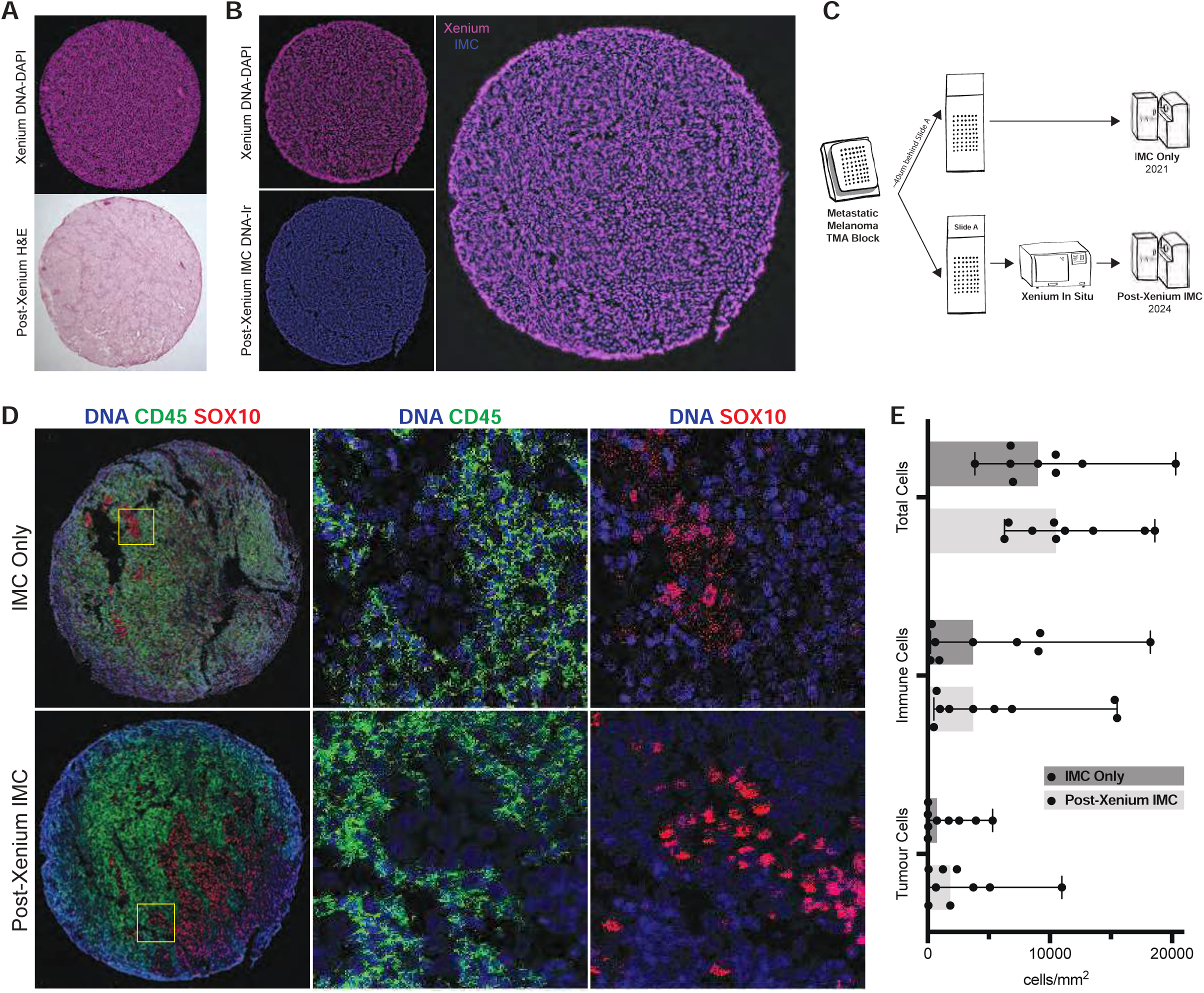
Xenium spatial transcriptomics has minimal impact on tissue integrity and subsequent quality of protein data generated by IMC. **A.** Top panel shows the Xenium DNA-DAPI image of a single tissue core (Slide B) at the start of the Xenium Analyzer workflow. Bottom panel shows H&E staining performed after Xenium on the same tissue core. **B.** Top-left panel shows the Xenium DNA-DAPI image of a single tissue core (Slide A). Bottom-left panel shows the corresponding IMC DNA-Ir image acquired on the same tissue core. Right panel displays the overlap of the Xenium DNA-DAPI image (magenta) and IMC DNA-Ir image (blue). **C.** Schematic overview of the two IMC experiments; ‘IMC Only’ and ‘Post-Xenium IMC’, performed on adjacent tissue sections from the same TMA tissue block. **D.** IMC images from adjacent tissue sections used in ‘IMC Only’ and ‘Post-Xenium IMC’ showing DNA (blue), CD45 (green) and SOX10 (red) staining of the same sample. **E.** Total nucleated cell, immune cell, and tumour cell densities for n=9 samples imaged in both the IMC Only (dark grey) and Post-Xenium IMC (light grey) experiments.

Next, we assessed the impact of the Xenium protocol on the quality of subsequent protein data from IMC. We compared the results of the IMC performed after Xenium (‘Post-Xenium IMC’, Slide A) to that of a separate IMC experiment (‘IMC Only’) that was performed alone on an adjacent tissue section from Slide A and B and acquired on the Hyperion Imaging System three years prior (Figure 2C). A comparison of the IMC images across the two experiments revealed that the Xenium protocol had little to no negative impact on subsequent IMC protein staining, highlighted here by comparable SOX10 staining for tumour cells and CD45 staining for immune cells (Figure 2D). Cellular quantification of nine samples from each IMC experiment confirmed that IMC performed after Xenium can be accurately analysed, as characterised by the protein signal quality generated for adequate detection of nucleated cells, tumour cells and immune cells that sits within the range of regular IMC experiments (Figure 2E).

### 3.3. Differences between Xenium and IMC datasets are apparent at the single-cell level

Next, we determined the level of concordance between the Xenium and IMC datasets in capturing key immune cell markers. First, we compared the spatial distribution of corresponding transcript and protein pairs imaged by the two technologies including the T cell receptor complex protein CD3E (*CD3E*) expressed by all T cells, CD20 (*MS4A1*) expressed by most B cells, and the cell proliferation marker Ki67 (*MKI67*) (Figure 3A). We observed a high visual correlation between the expression of proteins and corresponding transcripts for CD3E, CD20 and Ki67 at a global scale across samples, further validating the feasibility of performing IMC after Xenium on the same tissue section. However, when we compare the proteomic and transcriptomic datasets at a single-cell level, differences become apparent. This is evident in the disparity between the number of cells within a sample that are positive by transcript compared to protein for the markers CD3E, CD20 and Ki67, as demonstrated by both proportion and density (Figure 3B, C).

**Figure 3.**
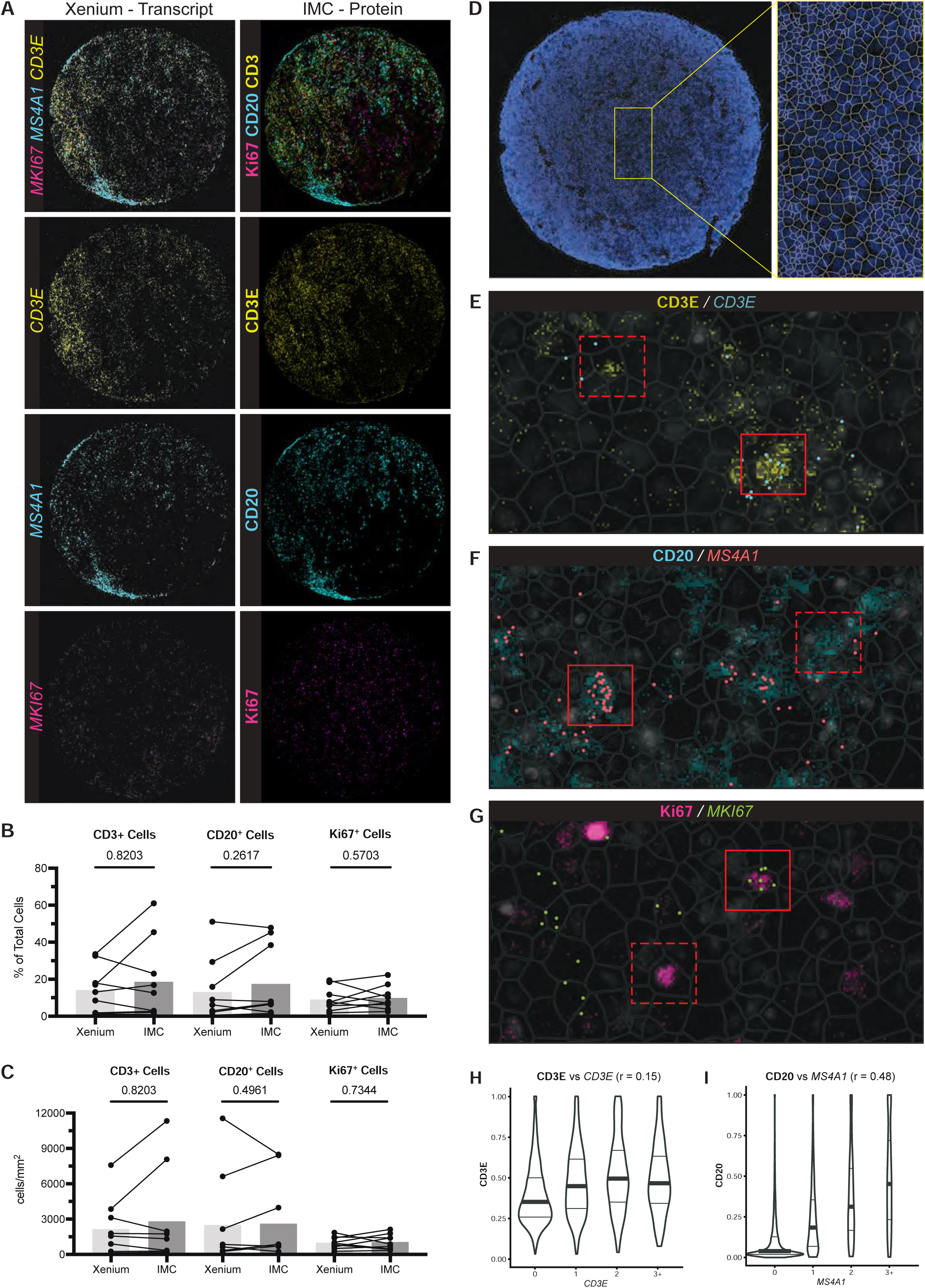
Differences between the expression of RNA transcripts (Xenium) and proteins (IMC) become apparent at the single-cell level. **A.** Left column shows the expression of the transcripts *CD3E* (yellow), *MS4A1* (cyan) and *MKI67* (magenta) for a single tissue core imaged by Xenium, each point represents one transcript. Right column shows the expression of the corresponding proteins CD3E (yellow), CD20 (cyan) and Ki67 (magenta) from the same tissue core, analysed by IMC. The proportion (**B**) and density (**C**) of cells in n=9 tissue cores positive for the protein markers CD3E, CD20 and Ki67 measured by IMC, compared to the corresponding transcripts measured by Xenium. Wilcoxon matched-pairs signed rank test was used. **D.** Co-registered IMC DNA-Ir image of a single tissue sample overlayed with the Xenium cell segmentation mask (zoomed-in image of the overlay) at the single-cell level. **E-G.** Zoomed-in images from the same tissue core showing the overlap of protein and transcript pairs CD3E (yellow)/*CD3E* (cyan) (E), CD20 (cyan)/*MS4A1* (red) (F) and Ki67 (magenta)/*MKI67* (lime) (G) at the single-cell level. Each image includes four layers: DNA-DAPI nuclei image (light grey) from Xenium, Xenium cell segmentation mask (dark grey outline), transcript points and corresponding protein stain. Solid-line box highlights examples of cells that are positive by both transcript and protein, whilst the dashed-line box indicates cells that are positive by transcript or protein alone. **H-I.** Correlation between the expression level of proteins (y-axis) and number of transcripts (x-axis) for CD3E/*CD3E* (H) and CD20/*MS4A1* (I) in a single tissue core.

The differences between Xenium transcript and IMC protein expression is also apparent after image registration. The Xenium and IMC datasets were integrated by first registering the corresponding nuclei images in Visiopharm. The Xenium cell segmentation mask generated onboard the Xenium Analyzer was subsequently applied to the co-registered images to generate a cell-by-feature matrix, where each cell had expression values for 377 transcripts and 40 protein markers (Figure 3D). Although we can observe clear instances where cells are positive by both transcript and corresponding protein for most marker pairs in the aligned images, there are also cells that have clear protein staining without transcript detected, and vice versa (Figure 3E-G). Next, we determined the correlation of corresponding marker pairs between platforms at the single-cell level. Some markers showed stronger correlation than others, in particular CD20*/MS4A1* (r = 0.48) compared to CD3E/*CD3E* (r = 0.15) (Figure 3H,I). Together our data demonstrates that while there is large similarity between transcripts and protein expression at a global scale, there is clear variability in concordance at the single-cell level, and therefore cell identities based on a single platform may have limitations.

### 3.4 Integrating transcriptomic and proteomic cell annotation provides a meaningful approach to understanding the complex nature of cellular landscapes

Cellular identity within tissues is fundamental to accurate interpretation of subsequent spatial analysis. We reasoned that integrated RNA and protein data could improve the accuracy of cell identities and provide greater clarity for mapping cellular interactions within fixed tissues. Rather than using a single transcript and single protein, we first determined whether multi-parameter identification of cell phenotypes in each method alone could improve concordance between the two platforms. This involved clustering the cells on co-registered Xenium and IMC datasets separately for manual cell annotation (Figure 3A, B). In the Xenium data, B cells were annotated based on the differentially expressed transcripts *MS4A1* and *CD79A,* whilst two T cell populations were identified by *CD3E* and either *CD4* or *CD8A* expression (Figure 4A). Similarly for the IMC data, B cells were identified by increased CD20 and HLA-DR protein expression, and T cells by increased CD3 and either CD4 or CD8A (Figure 4B). Once T cells and B cells were identified, we assessed the concordance between the two platforms when cells were identified solely based on the presence or absence of CD3E (*CD3E*) and CD20 (*MS4A1*) (1^st^ Approach), compared to the semi-supervised clustering approach based on multiple markers (2^nd^ Approach) (Figure 4C, D). Although total T cells identified increased and total B cells decreased in the clustering (2^nd^) approach (Figure 4E, F), it did improve the proportion of cells identified by both Xenium and IMC for both cell types (Figure 4G,H). Together, our data shows that multi-marker classification using semi-supervised clustering improves concordance of cell phenotypes between the platforms.

**Figure 4.**
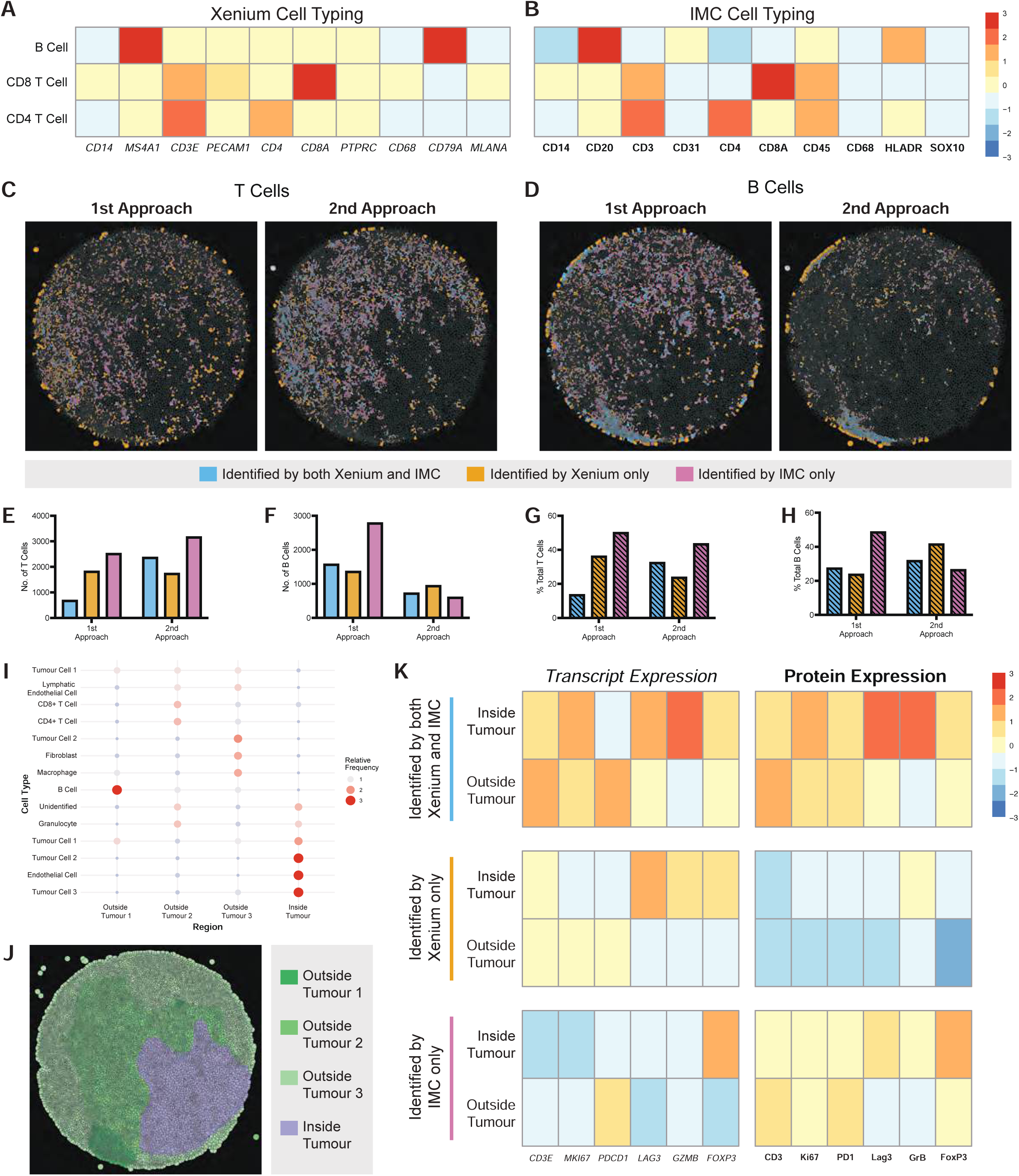
Integrating Xenium transcriptomics and IMC proteomics for more meaningful cell annotation and characterisation of tissue location-associated cell properties. **A.** clustSIGNAL clustering of the Xenium data was performed to identify 9 cell clusters. Here one B cell cluster and two T cell clusters, identified by manual cell typing, are represented by a heatmap of relative RNA transcript marker expression where ‘0’ represents average marker expression. **B.** FuseSOM clustering of the IMC data was performed to identify 10 cell clusters. Here one B cell cluster and two T cell clusters, identified by manual cell typing, are represented by a heatmap of relative protein marker expression. **C-D.** Concordance masks of annotated T cells (C) and B cells (D) identified by both Xenium and IMC (blue), Xenium only (orange), and IMC only (pink) in a single tissue sample. ‘1^st^ Approach’ represents the single marker approach for cell annotation, ‘2^nd^ Approach’ represents the semi-supervised clustering approach. **E-F.** The number of T cells (E) and B cells (F) identified by both Xenium and IMC (blue), Xenium only (orange), and IMC only (pink) for each approach in the single tissue sample. **G-H.** The proportion of total T cells (G) and B cells (H) identified by both Xenium and IMC (blue), Xenium only (orange), and IMC only (pink) for each approach in the single tissue sample. **I.** 4 regions in the single tissue sample were identified and manually annotated based on the relative frequency of cell types. **J.** Visual representation of the 4 regions in the single tissue sample, highlighting the single ‘inside tumour’ region (purple) and three distinct ‘outside tumour’ regions (green). **K.** Heatmap displaying the average expression of RNA transcripts and corresponding proteins identified by both technologies or by a single modality alone on T cells, further stratified by whether they were located inside or outside the tumour tissue region.

Next, we investigated the biological relevance of measuring both RNA and protein in spatial analysis by determining whether location-based differences could be identified when comparing T cells residing within tumours to their counterparts outside the tumours. This involved firstly annotating ‘inside tumour’ and ‘outside tumour’ regions based on the relative frequency of immune and tumour cell types within each identified region (Figure 4I, J). As expected, T cells within tumours were associated with increased activation, proliferation and effector functions compared to T cells outside the tumours (Figure 4K). T cells identified by both platforms had increased expression of the transcripts *MKI67*, *LAG3*, *GZMB* and *FOXP3* in tumour regions relative to non-tumour regions, with expression of the corresponding proteins identified at the protein level. In contrast, but not surprisingly, T cells identified by Xenium alone mainly showed increased expression of select markers at the transcript but not the protein level, whilst T cells identified by IMC alone primarily showed increased expression at the protein level. Together, our data shows that cell populations identified using markers detected by both platforms produces the most biologically accurate results at both the transcript and protein level. Therefore, integrating transcriptomic and proteomic cell annotation by combining Xenium and IMC provides a meaningful method for the in-depth characterisation of cell phenotypes across spatial locations, and could be used to further understanding of functional capabilities and cellular interactions.

## 4.0 Discussion

Although many spatial technologies currently available can provide an expansive view of tissue immune microenvironments, most platforms are limited to assessment of RNA transcripts or protein profiling alone. Whilst protein expression is more dependable for determining cellular phenotypes, it has restricted profiling capabilities compared to transcriptomics and could limit the molecular characterisation of different immune subsets. Here, we aimed to explore the potential of combining two single-omic spatial modalities to achieve a more comprehensive view of cellular phenotypes and landscapes. We demonstrate the feasibility of performing Xenium and IMC on a single FFPE tissue section, enabling the spatial mapping of both high-parameter transcriptomics and targeted proteomic analysis at single-cell resolution. In accordance with other dual-platform workflows (Huynh et al., 2025; Rademacher et al., 2025; Tran et al., 2025), we show that the integrity of the tissue is largely preserved after the cyclic Xenium protocol and is suitable for subsequent protein staining. Additionally, we confirm that IMC signal intensity is well maintained after Xenium staining, further validating the practicality of the workflow.

In line with previous studies (Cervilla et al., 2025; Huynh et al., 2025; Tran et al., 2025), our work highlights the differences present between transcript and protein expression in immune cells in FFPE tissue. This discordance is to be expected, as biologically there are multiple reasons why transcript expression may not always correlate with protein, including post-transcriptional and post-translational modifications, protein half-lives, and cellular conditions (Buccitelli and Selbach, 2020). Thus, this workflow can additionally be used to provide further insights into how protein production and expression is regulated within immune cells. Interestingly, there is evidence to suggest that some immune markers are more correlated at the transcript and protein level than others. A study analysing PBMCs by CITE-seq found that CD20 along with CD8a, CD11b and CD16 correlated relatively well with their corresponding transcript, whilst other markers including CD3, CD4 and CD69 showed greater disagreement (Li et al., 2020). This mirrors findings from our work, where we found CD20 had a better RNA and protein correlation than CD3. These differences are also apparent in two dual-platform spatial studies combining Xenium with either COMET or PCF, with both similarly observing relatively low correlations between transcript and protein expression for immune markers (Huynh et al., 2025; Tran et al., 2025). However, it is also possible that the discordance between RNA and protein seen in our study and others are due to experimental and technical challenges, including poor segmentation, poor registration, and RNA diffusion. Altogether, our work combined with previous findings emphasises how relying on transcript or protein expression alone may lead to an inaccurate representation of cell populations in tissues, thus highlighting the importance of combining the two modalities.

The value of integrated transcriptomic and proteomic data cannot be overstated. Current spatial proteomic workflows, including IMC, rely on a targeted approach, and the use of antibodies limits the number of parameters that could be identified in each cell population. This is particularly challenging when various subsets of immune cells are defined by multiple markers. Transcriptomic data alone also has major limitations, including instances of off target binding with the Xenium platform (Hallinan et al., 2025) and marker contamination, which could occur due to poor segmentation, RNA diffusion, and overlapping cells. Importantly, defining the nature of individual cell-to-cell interactions in tissues requires accurate identification of cell types and the molecular signalling pathways that are associated with them. Supplementing protein spatial analysis with same-slide transcriptomics by integrating Xenium and IMC is a promising approach to achieve this, as illustrated here where we showed that T cells identified by both platforms had the most biologically meaningful location-based differences at both the transcript and protein level, as opposed to cells identified by a single platform. This combined approach clearly demonstrates that T cells inside tumour regions have different functional potential compared to their counterparts outside tumour regions. This is reflected by increased expression of activation, proliferation and effector function markers which align with T cell capabilities such as cytotoxic killing of tumour cells. Together, our data shows that cell populations identified by combining multi-modal characteristics of cells in tissues provides a meaningful approach to understanding the nature of complex cellular interactions.

There are several challenges to be considered when integrating spatial transcriptomics and proteomics data from the same tissue section. This includes a potential decrease in the quality of the secondary dataset acquired, that being IMC in this workflow. However, in our experimental workflow, we did not observe a decline in protein (IMC) signal intensity or tissue integrity on the processed Xenium slide. Despite this, other experimental limitations may have impacted the quality of the IMC data and contributed to the differences in protein and RNA expression observed between the two platforms. These limitations included an extended time between Xenium analysis and IMC staining, in this study IMC staining was performed nine days after the conclusion of the Xenium run, which is outside the 1-week storage time suggested by 10X Genomics for post-Xenium experiments. Secondly, as this workflow was performed on FFPE samples collected between 2001 and 2007, there is also the possibility of both RNA and protein degradation over time (Hester et al., 2016; Wolff et al., 2011). It is also possible that reagents used in the Xenium staining may have impacted these samples, despite limited evidence of this in our study, as proteases such as those used in the protocol are known to impact tissue morphology (Chiecchio, 2020; Lemes et al., 2024; Selvarajan et al., 2003).

Alongside experimental limitations, there are also technical hurdles concerning registration that must be considered in the integration of this data. Registration of the two data sets poses a challenge due to fundamental differences in how the data is acquired by the separate platforms, with Xenium acquiring data as images taken in consecutive cycles and IMC data acquiring all signals simultaneously per um^2^ of tissue ablated. Although both platforms ultimately produce images at the single-cell level, the respective nuclei images are not identical. This is likely due to the different methods of DNA identification Iridium Intercalator (IMC) compared to DAPI (Xenium), as well as possible tissue shrinkage after IMC staining due to samples being air-dried before ablation in the Hyperion instrument. Additionally, tissue artefacts that can occur throughout the experimental or imaging process, causing focusing issues during Xenium image acquisition could cause further local and global differences between the images acquired. To account for this, we manually registered the two datasets by placing landmarks on corresponding nuclei in the Xenium DNA-DAPI and IMC DNA-Ir images. Despite achieving sufficient registration with Visiopharm software in this study, future work will involve exploring computationally driven registration approaches to increase sample throughput. Ideally, registration would be based on images both derived from the same stains such as DAPI or a cell-membrane stain, as is the case for the dual-platform workflows combining Xenium with the fluorescent-based proteomic platforms COMET and PCF. Despite these challenges, here we were still able to demonstrate the successful registration of Xenium and IMC cell and tissue data for the first time.

Cell segmentation is an additional challenge in integrated multi-omic spatial datasets. For our analysis, we chose to use the Xenium Analyser cell segmentation mask as opposed to the IMC mask generated using CellProfiler software for analysis of the aligned datasets in this study. After inspecting both masks, it was determined that the Xenium mask provided a more accurate segmentation of the cell landscape. Additionally, Xenium automatically generates a cell segmentation mask, in contrast to CellProfiler and many other segmentation techniques which require extensive manual adjustment of segmentation parameters. In spatial biology, cell segmentation remains an area where there is room for improvement. Xenium explorer masks are individually created by nuclear expansion of 15 um or until another cell boundary is encountered, which is not always biologically accurate due to differences in cell morphologies and anucleated cells and structures in tissues. Recently improved Xenium assays now offer a cell segmentation kit that includes antibodies that bind to the cell boundary and intracellular components in combination with a promising multimodal algorithm for more accurate cell segmentation. Future work will assess improved computational and experimental cell segmentation methods.

Here we have described the novel dual-platform workflow that combines Xenium and IMC on a single FFPE tissue section to facilitate spatial profiling of both mRNA and protein at single cell resolution. This multi-omic method has the potential to contribute to more accurate and meaningful characterisation of cell populations, their interactions within their spatial locations and their functional potential. Importantly, this provides a more comprehensive view of tissue microenvironments for furthering our understanding of complex biological disease processes.

## Acknowledgements

This project was performed with funding received from SURFebruary grant from the Chris O’Brien Lifehouse. ROA was funded by an Australian Government Department of Education RTP scholarship & the Melanoma Institute of Australia Postgraduate Research Top-Up Scholarship. XB is supported by the University of Sydney and Melanoma Institute Australia Scholarships. Analysis was performed at the Sydney Cytometry Core Research Facility at the Charles Perkins Centre within the University of Sydney.

